# *In vitro*-generated inflammatory fibroblasts secrete extracellular matrix with biochemical and biophysical properties similar to tissue-remodelling fibroblasts

**DOI:** 10.1101/2024.09.26.614950

**Authors:** Fàtima de la Jara Ortiz, Chiara Cimmino, Kurt Grech, Martijn A. Huynen, Eline Janssen, Vera Wagenaar, Maxime C. van Zwam, Koen van den Dries, Maurizio Ventre, Alessandra Cambi

## Abstract

In solid cancers, inflammation and viral infections, two main fibroblast subtypes have been identified: myofibroblast-like fibroblasts and inflammatory fibroblasts. In the tumour microenvironment (TME), these cancer-associated fibroblast (CAF) subtypes are known as myCAFs, which generate a stiffened fibrotic extracellular matrix (ECM), and iCAFs, which secrete inflammatory cytokines to locally modulate the immune response. Yet, whether iCAFs contribute to shaping the ECM biochemical and biophysical properties remains unknown, mainly because robust *in vitro* models to generate fibroblast subtypes are lacking. Here, we established an *in vitro* cell culture system based on murine NIH3T3 fibroblasts and stimulation by TGFβ and IL1α, alone or in combination, to induce fibroblast subtypes. Gene expression analysis of well-documented myCAF (*Acta2*/*Tagln*) and iCAF (*Ccl2*/*Il6*/*Lif*) markers revealed that TGFβ induced a myCAF-like phenotype, while a combination of TGFβ and IL1α induced an iCAF-like phenotype. We compared these *in vitro* subtypes to myCAFs and iCAFs from publicly available scRNAseq data of tumour tissues from cancer patients. We found that, similar to myCAFs, both tumour-associated and *in vitro* iCAFs express *Acta2*/*Tagln* as well as genes encoding for typical ECM proteins, which correlated *in vitro* with the ability to contract collagen. Furthermore, fluorescence microscopy and atomic force microscopy revealed that *in vitro* both subtypes generate thick, layered and stiff matrices with highly aligned ECM, demonstrating for the first time that iCAFs may also contribute to a pathological ECM. Finally, matrices generated from these *in vitro* fibroblast subtypes, but not from uninduced or IL1α-only stimulated fibroblasts, enhanced the expression of the immune suppression marker Arg1 in co-cultured macrophages. Our study provides new insights in the contribution of inflammatory fibroblasts to ECM deposition and remodelling and puts forward a well-defined *in vitro* model to generate different fibroblast subtypes for future in-depth mechanistic studies of their roles in cancer and other pathologies.

## Introduction

Fibroblasts are classically considered the major producers and remodellers of the extracellular matrix (ECM) both in tissue homeostasis and pathological conditions. In several diseases such as infection, inflammation and cancer, fibroblast populations can expand and acquire pathological phenotypes driving disease progression and influencing therapy response (1–4). Recent studies in arthritis, viral infection and cancer have revealed that, besides the ECM-generating fibroblasts, other fibroblast subtypes exist with immune modulatory properties (3, 5–7). Whether differences in ECM modulatory properties exist between these fibroblast subtypes is, however, largely unexplored.

Fibroblasts are a major cellular components of the tumour microenvironment (TME) in solid tumours (8–14). They generate, deposit and remodel the tumour ECM, which is a key determinants of tumour cell behaviour and disease outcome (13–16). ECM biophysical properties such as stiffness or alignment are also critical for controlling cancer cell migration (17, 18), proliferation (19–21) and differentiation (19–22). Furthermore, aberrant ECM composition and architecture have been associated with therapy resistance and poor prognosis (23, 24). Numerous studies to better understand the crucial role of fibroblasts in shaping the TME ECM have led to the identification of various cancer-associated fibroblast (CAF) subtypes (25). Specifically, single-cell RNA sequencing (scRNAseq) of stromal cells from patient samples has recently led to the identification of two main subtypes, myofibroblast-like CAFs (myCAFs) and the inflammatory CAFs (iCAFs), characterized by the expression of *Acta2*/*Tagln* and *Ccl2*/*Il6*/*Lif*, respectively (26–30).

Functionally, myCAFs have been described to be primarily responsible for ECM production and deposition, and present a contractile phenotype (6, 31). Furthermore, they share many features with fibrosis-associated myofibroblasts, which are known to contribute to a stiffened fibrotic ECM (31). iCAFs, on the other hand, are reported to have a secretory phenotype, releasing inflammatory cytokines and contributing to shape the immune state of the TME (6, 32). iCAFs, however, have been described to express genes related to collagen production (33), ECM deposition and organization in pancreatic (29) and bladder (34) tumours. Yet, their ability to shape the biochemical and biophysical properties of the ECM remains unknown.

*In vivo* and *in vitro* models both in human and mouse have been established to classify and further interrogate arising CAF subtypes. Most *in vivo* animal models allow for CAF subtype identification and offer opportunities to dissect functional interactions between CAFs and other cell types within the TME (26, 27, 29, 30, 32, 35–37). Yet, animal models are bound to the tumour type and allow limited cellular manipulation (38). *In vitro* models, on the other hand, are more standardized and versatile, allowing controlled manipulation of fibroblasts and their interaction with cancer cells, but lack the pathophysiological complexity of the TME (6, 32, 39). myCAFs and iCAFs have been successfully modelled *in vitro* using primary cells from pancreatic ductal adenocarcinoma (PDAC), serving as a ground-breaking tool to study subtype-specific characteristics as well as intra-tumour CAF heterogeneity and plasticity (6, 32). In these studies aiming at understanding how tumours affect TME resident cells, conditioned medium (CM) from tumour cells has often been used to induce CAF differentiation (6, 32, 40). Despite being a powerful tool, the composition of tumour cell CM can vary depending on tumour type or culturing conditions, which can be a source of variability among different studies. Mimicking the induction of myCAFs and iCAFs in PDAC using primary pancreatic stellate cells and KPC tumour organoids has confirmed that *in vitro* myCAFs are TGFβ-inducible, while IL1α promotes iCAF induction (32). The interplay between TGFβ and IL1α is of interest as their pathways are known to harbour a reciprocal regulation, either through SMAD7 or TAK1 (32, 41–44). Since in the TME, both TGFβ and IL1α are likely simultaneously present, investigating their mutual effects on shaping the composition and plasticity of the CAF population is warranted.

Besides cancer, fibroblast subtypes with phenotypes that are quite similar to myCAFs or iCAFs are also present in other pathologies. Hence, there is a clear need for robust, standardized and reproducible *in vitro* models of fibroblast subtype generation that allow detailed mechanistic studies of subtype-specific functions and their relation to the biophysical properties of pathological ECMs.

Here, we established a well-defined *in vitro* cell culture system based on murine NIH3T3 fibroblasts and stimulation by TGFβ and IL1α, alone or in combination, to induce fibroblast subtypes. As substantial knowledge about tumour-resident fibroblast is available, we benchmarked our *in vitro* fibroblast subtypes against myCAFs and iCAFs that have been previously defined by scRNAseq. Our results demonstrate that myCAF-like fibroblasts can be induced by TGFβ and revealed that both TGFβ and IL1α are needed to induce iCAF-like fibroblasts. We provide a detailed characterization and comparison of the composition, architecture and biophysical properties of the ECMs generated by these *in vitro* subtypes. Lastly, we interrogated the capability of the fibroblast subtype-specific ECMs to influence macrophage polarization. Our study provides new insights on the biology of inflammatory, iCAF-like fibroblasts revealing their potential contribution to pathological ECM composition and biophysical properties. This well-characterized *in vitro* model to generate tissue-remodelling and inflammatory fibroblasts can be used for future in-depth mechanistic studies of the TME or other pathology in which fibroblast subtypes are involved.

## Results

### *In vitro* induction of fibroblast subtypes that are phenotypically similar to myCAFs and iCAFs of tumour tissues

In order to develop a robust and well-controlled *in vitro* model of myCAF and iCAF subtypes, we cultured NIH3T3 fibroblasts in a 3D fibrillar collagen gel for 4 days in the presence of either TGFβ or IL1α, which have previously been described to induce myCAFs and iCAFs, respectively, when added to pancreatic stellate cells (6, 32). As TGFβ and IL1α are likely often present at the same time in the TME, we also stimulated fibroblasts simultaneously with TGFβ and IL1α. In all experiments, fibroblasts cultured with medium only (uninduced) were included as control (**Fig. 1A**). After 4 days of culture, we assessed the gene expression of conventional myCAF markers (*Acta2* and *Tagln*) and iCAF markers (*Ccl2, Il6* and *Lif*) (26–30). We found that, as compared to uninduced fibroblasts, both *Acta2* and *Tagln* expression was higher in TGFβ-stimulated fibroblasts, while it was lower in IL1α-only stimulated fibroblasts (**Fig. 1B**). When stimulated with both TGFβ and IL1α, *Acta2* and *Tagln* levels were similar or higher as compared to uninduced fibroblasts (**Fig. 1B**). For the iCAF markers, we observed a high expression of *Ccl2, Il6* and *Lif* when IL1α is present, and no or only nonsignificant increases when only TGFβ was added to the cell culture. Interestingly, the strongest increase in iCAF marker expression was observed when fibroblasts were stimulated with the combination of TGFβ and IL1α, suggesting an additive, perhaps synergistic effect (**Fig. 1C**).

**Fig. 1.**
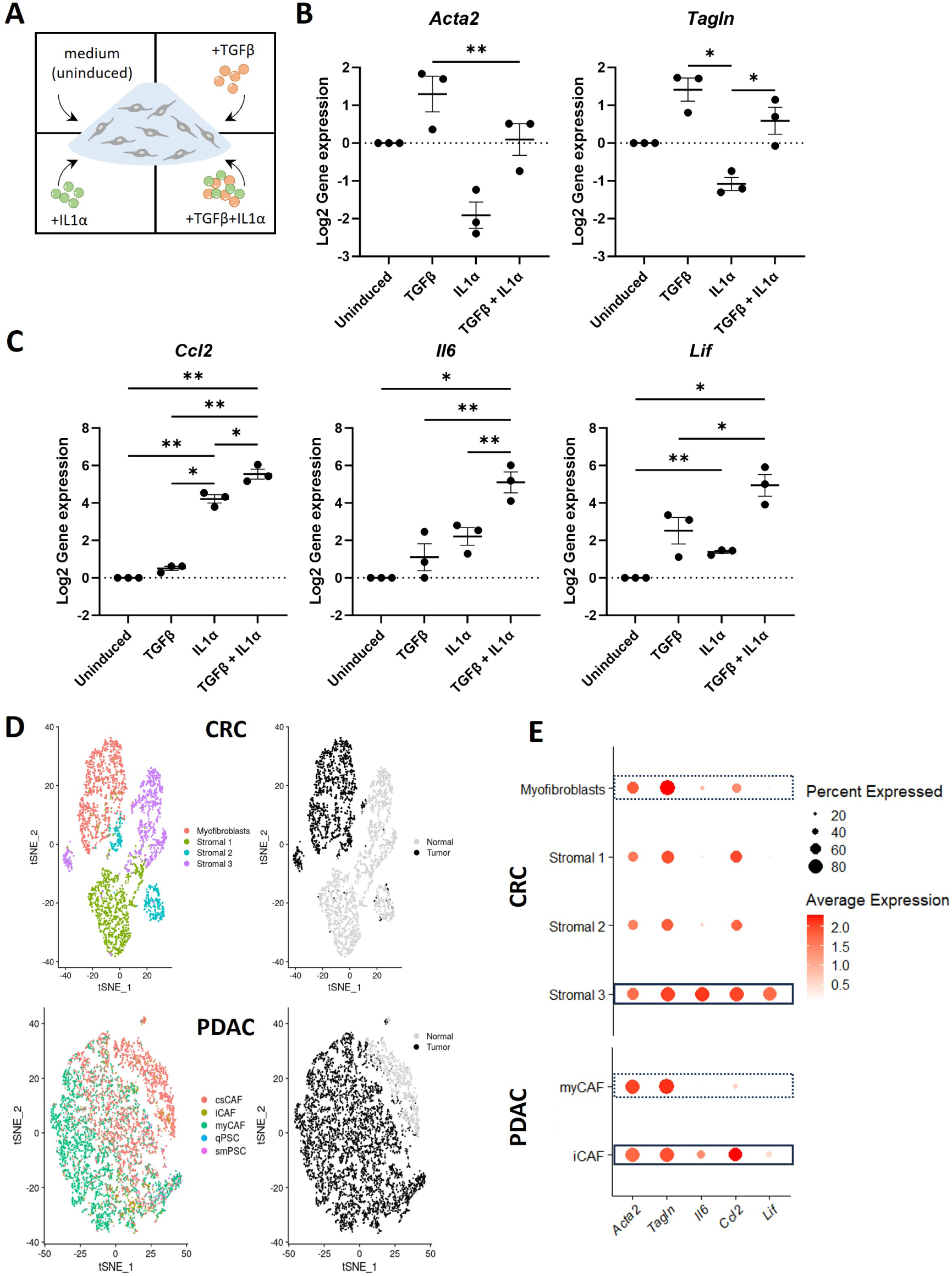
*In vitro* generation of fibroblast subtypes. **A)** Schematic representation of the stimuli used to induce fibroblast differentiation *in vitro* in 3D. NIH3T3 fibroblasts were mixed with a drop of rat tail fibrillar collagen I (1.7mg/mL polymerized at 37°C) and cultured for 4 days in the presence of medium alone or medium supplemented with the indicated stimuli. **B-C)** Log2 gene expression of myCAF markers *Acta2* and *Tagln* (B) and *Ccl2, Il6* and *Lif* (C) of *in vitro*-induced subtypes normalized to uninduced. Housekeeping genes are *Ppia* and *B2m*. Data were analysed by one-way ANOVA followed by Tukey’s multiple comparisons test of three independent experiments. *p<0.05; **p<0.01. **D)** t-SNE plots of 3,392 stromal fibroblasts of interest from CRC dataset (top) and PDAC dataset (bottom), color-coded by sub-cell type and sample origin, and 6,781 stromal fibroblasts of interest. **E)** Dot-plots representing average gene expression, and percentage of cells expressing *Acta2, Tagln, Il6, Ccl2* and *Lif*. From the CRC datasets (top), a comparison is made between subtypes Myofibroblasts, Stromal 1, Stromal 2, and Stromal 3 of tumor origin. From the PDAC dataset (bottom), myCAFs and iCAFs are compared.

To determine to what extent our *in vitro* generated fibroblast subtypes relate to tumour-resident myCAFs and iCAFs, we reanalysed publicly available scRNAseq datasets of primary colorectal cancer (CRC) (36) and pancreatic ductal adenocarcinoma (PDAC) (37) from patients and plotted the expression of myCAF (*Acta2, Tagln*) and iCAF markers (*Il6, Ccl2, Lif*) of the CAF populations identified in the tumour tissues (**Fig. 1D, E**). For clarity, while the CRC dataset identifies myofibroblasts and stromal 1, 2 and 3 populations as fibroblasts present in tumour tissue, the PDAC dataset does not classify tumour fibroblasts into different subpopulations. We therefore used the validated transcriptomic atlas of the PDAC TME (45) to classify the fibroblast subpopulations in the PDAC dataset.

Then we re-clustered the fibroblasts and retrieved the various fibroblast types that were originally identified as either tumour-resident or normal tissue-resident and displayed them using tSNE (**Fig. 1D**). As expected, *Il6, Ccl2* and *Lif* were highly expressed by the iCAFs (i.e. stromal 3 population) of the CRC dataset and in the iCAF population of the PDAC dataset (**Fig. 1E**). Yet, the expression of the classical myCAF markers *Acta2* and *Tagln* was not restricted to myCAFs but was clearly detected at similar levels also in the iCAFs of both tumour types (**Fig. 1E**). When correlating these findings to our *in vitro* culture system, we concluded that the TGFβ-stimulated fibroblasts resemble *in vivo* myCAFs, while the fibroblasts stimulated with both TGFβ and IL1α resemble the tumour-associated iCAFs.

In the remainder of our manuscript, we will therefore refer to the TGFβ-stimulated fibroblasts as *in vitro* myCAFs with tissue-remodelling capacity and to the TGFβ+IL1αstimulated fibroblasts as *in vitro* iCAFs with inflammatory phenotype. Importantly, in our experiments we will continue to include the IL1α-only stimulated and the uninduced fibroblasts for comparison.

### *In vitro* myCAFs and iCAFs have similar capacity to contract the surrounding matrix

myCAFs are considered the fibroblasts specialized in ECM production and remodelling due to their highly contractile capacity, but less is known regarding iCAFs (6, 39, 46, 47). To functionally assess this aspect, we designed a floating collagen ring contraction assay (**Fig. 2A**). In this assay, a freshly polymerized collagen ring containing fibroblasts is released from the culture dish and allowed to float in culture medium. Over time, the ring is contracted due to the fibroblast contractile activity, resulting in a smaller ring area. We prepared collagen rings containing fibroblasts induced with TGFβ or IL1α or a combination thereof as well as collagen rings containing uninduced fibroblasts in the absence or presence of the Rock inhibitor Y-27632 and measured the ring area after 2 days of culture (**Fig. 2B**). As Y-27632 fully inhibits cell contractility, we set this condition as the negative control and normalized all the ring areas to the area of these samples (**Fig. 2C**). The uninduced condition indicates the fibroblast intrinsic ability to contract, which was similar to the contraction observed for IL1α-induced fibroblasts. In contrast, both *in vitro* myCAFs and *in vitro* iCAFs show an evident and quite comparable contraction of the ring (**Fig. 2B,C**).

**Fig. 2.**
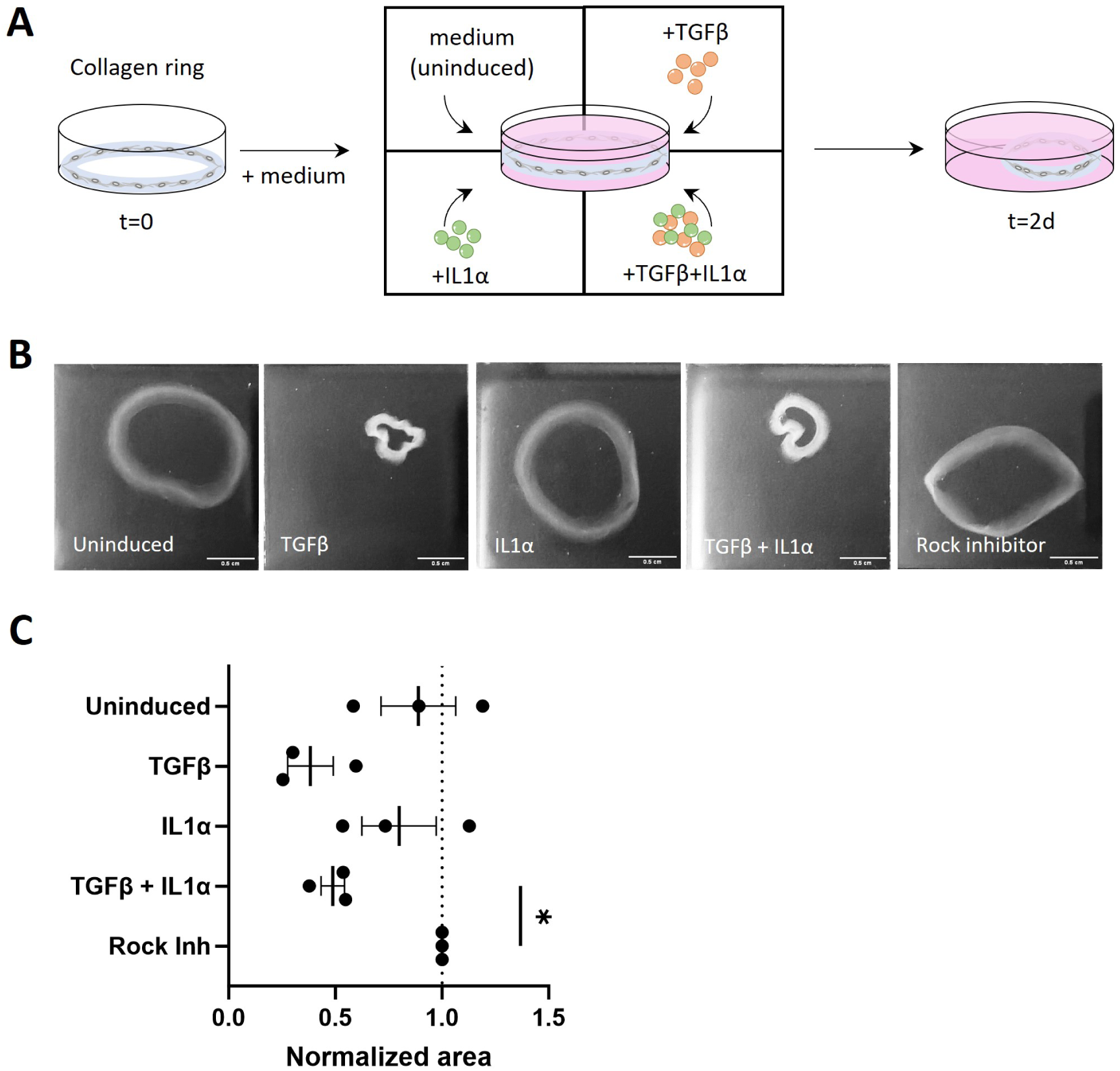
*In vitro* myCAFs and iCAFs have similar capacity to contract the ECM. **A)** Schematic representation of the collagen contraction assay. **B)** Images of collagen ring contraction under different conditions. Scale bar = 0.5 cm. **C)** Collagen ring contraction quantification normalized to the Rock Inhibitor condition that represents the negative control. Data were analysed by one-way ANOVA followed by Dunnett’s multiple comparisons test of three independent experiments. *p<0.05.

These results demonstrate that the high expression levels of *Acta2* and *Tagln* detected in both *in vitro* myCAFs and *in vitro* iCAFs indeed correlate with a highly contractile phenotype of these CAF subtypes.

### *In vitro* myCAFs and iCAFs produce layered cultures with similar cell organization

myCAFs are reportedly the major cell type responsible for the stroma formation in the TME, but little is known regarding iCAF contribution to the ECM composition and organization (13, 48). Considering the strong similarity we found between *in vitro* myCAFs and *in vitro* iCAFs in terms of contractility, we sought to investigate whether these *in vitro* CAF subtypes are equally capable of producing stromal-like matrices. For this, we established a 2 week-long 2D culture system that allowed the fibroblasts to generate subtype-specific cell derived matrices (CDMs), as reported in previous studies (49, 50) (**Suppl. Fig. 1A**). As addition of ascorbic acid is necessary to stimulate collagen production in cultured fibroblasts (49, 51), we first ascertained that this chemical did not alter the expression pattern of the fibroblast subtype-specific markers (**Suppl. Fig. 1B**). Also, to determine whether the induced phenotype was maintained over time, we evaluated the expression of the CAF subtype markers in fibroblasts isolated from the CDMs and confirmed *Tagln* expression in both *in vitro* CAF subtypes and *Ccl2* expression in the *in vitro* iCAFs, albeit at lower levels with respect to earlier time points (**Suppl. Fig. 1B**).

To subsequently determine the organization of the cells in these CDMs, we stained the samples for filamentous actin (F-actin) and the nuclei and imaged them by fluorescence microscopy. The different stimulations visibly affected the fibroblast morphology, which we quantified by measuring the directionality of the actin cytoskeleton and of the nuclei (**Fig. 3A, B**). In both uninduced and IL1α-only induced cultures, fibroblasts are randomly oriented and have round nuclei. Conversely, *in vitro* myCAFs and *in vitro* iCAFs appear more elongated with oblong nuclei, and are clearly coaligned forming hundreds of micron wide patches of ordered tissues (**Fig 3A, B**).

**Fig. 3.**
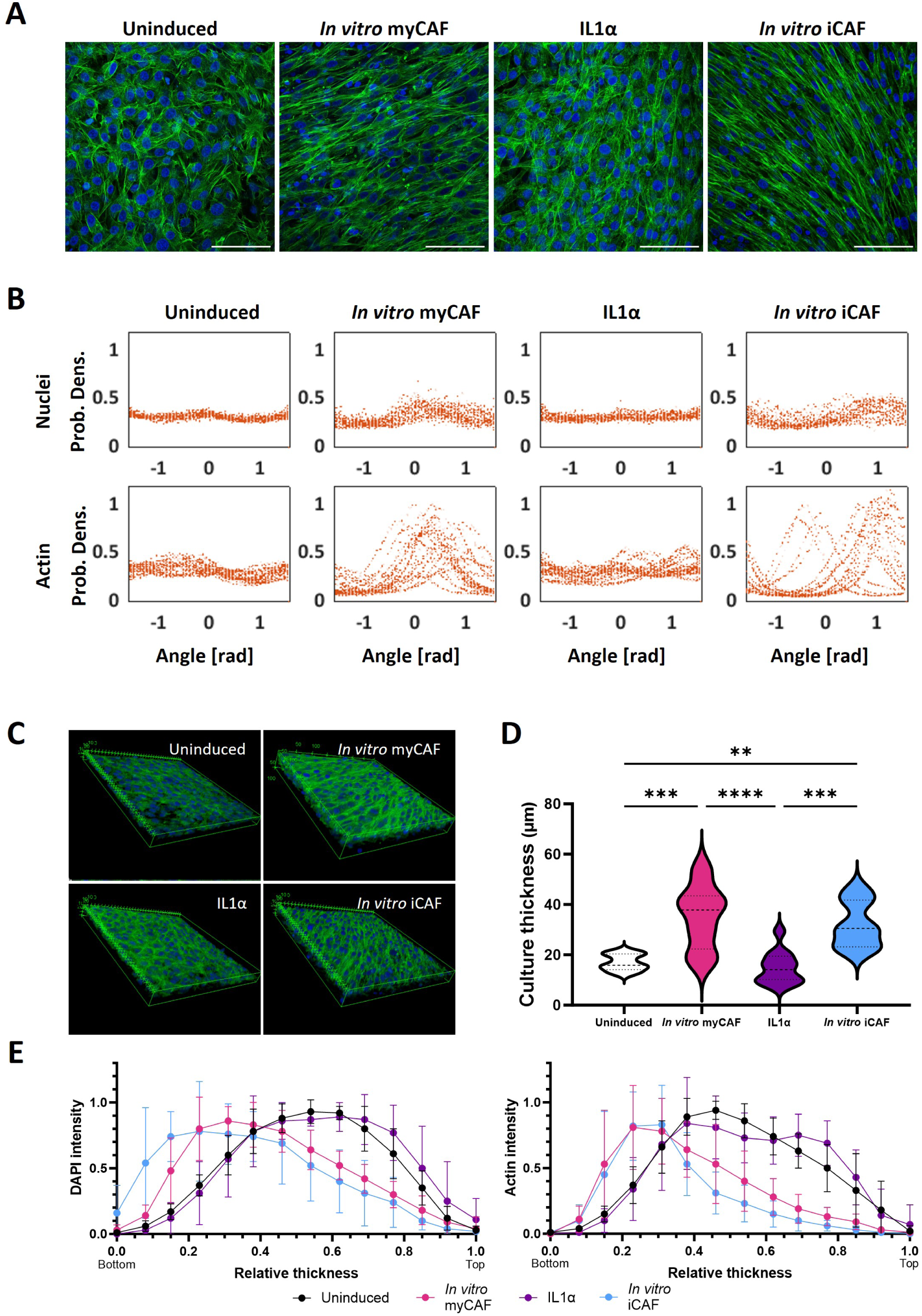
*In vitro* myCAFs and iCAFs produce layered matrices with similar cell organization. **A)** Immunofluorescent staining of nuclei (blue) and actin (green) of cells belonging to matrices induced in different conditions. Scale bar = 100um. **B)** Directionality of the cells is measured by “Directionality” plugin in FIJI. The resulting angle is a measurement from 0 to 1.5 or -1.5 radians. The probability density function represents the probability of observing the event in that particular angle. **C)** 3D reconstruction of subtype cultures from Z stack images of actin (green) and Dapi (blue) obtained by confocal microscopy (354.25x 354.25 μm). **D)** Thickness quantification of subtype cultures based on immunofluorescent actin signal. For uninduced and *in vitro* iCAF cultures, 9 measurements where performed, while 12 were performed for *in vitro* myCAF and IL1α. Data were analysed by one-way ANOVA followed by Tukey’s multiple comparisons test. **p<0.01; ***p<0.001; ****p<0.0001. **E)** Axial localisation of the fibroblasts in the layered matrices. Dapi and actin fluorescence intensity along the thickness of the matrices was normalized to the maximum intensity of each condition. Dots represent the mean ±SD of 6 samples for uninduced, *in vitro* myCAF and IL1α matrices, and 3 samples for *in vitro* iCAF cultures.

We then quantified the axial organization and thickness of these cultures and found that under all conditions cell layers were generated (**Fig. 3C-E, Suppl. Movie 1**). Yet, the CDMs generated by *in vitro* myCAFs and *in vitro* iCAFs are significantly thicker, as compared to those produced by uninduced or IL1α-induced fibroblasts (**Fig. 3C,D**). In addition, in these CDMs, cells mostly accumulated in the basal strata of the tissue, whereas in the CDMs from either uninduced or IL1α-only induced fibroblasts cells were more homogeneously distributed throughout the culture layers (**Fig. 3E**).

Collectively, these results indicate that *in vitro* myCAFs and *in vitro* iCAFs are able to generate thick stromal layers with similar cellular alignment and axial localization.

### *In vitro* myCAFs and iCAFs produce matrices of comparable composition and biophysical properties

The larger thickness and the characteristic accumulation of the fibroblasts in the most basal strata observed in *in vitro* myCAF and iCAF cultures compared to uninduced and IL1αonly suggest that these *in vitro* CAF subtypes could both produce ECM components, which are usually deposited on top of the cell layers. Gene expression analysis of the induced fibroblasts at day 4 interestingly showed similar *Col1a1* upregulation in both *in vitro* CAFs (**Suppl. Fig. 1B**). This prompted us to explore the expression levels of several genes involved in ECM production, deposition and remodelling in the tumour tissue iCAFs of the CRC and PDAC scRNAseq databases. We observed that *in vivo* tumour-associated iCAFs and myCAFs had similar levels of elastin, fibronectin and collagens as well as similar levels of collagen crosslinking enzymes such as the lysyl oxidases *Lox* and *Loxl1* (**Fig. 4A)**. Together, this database analysis shows that iCAFs express a set of genes involved in ECM production that is present in all the other CAF populations identified, strongly indicating that *in vivo* not only myCAFs but also iCAFs are able to produce and remodel the stromal ECM. The *in vitro* CAF subtypes therefore fairly resemble the *in vivo* myCAFs and iCAFs in terms of expression of ECM related genes.

**Fig. 4.**
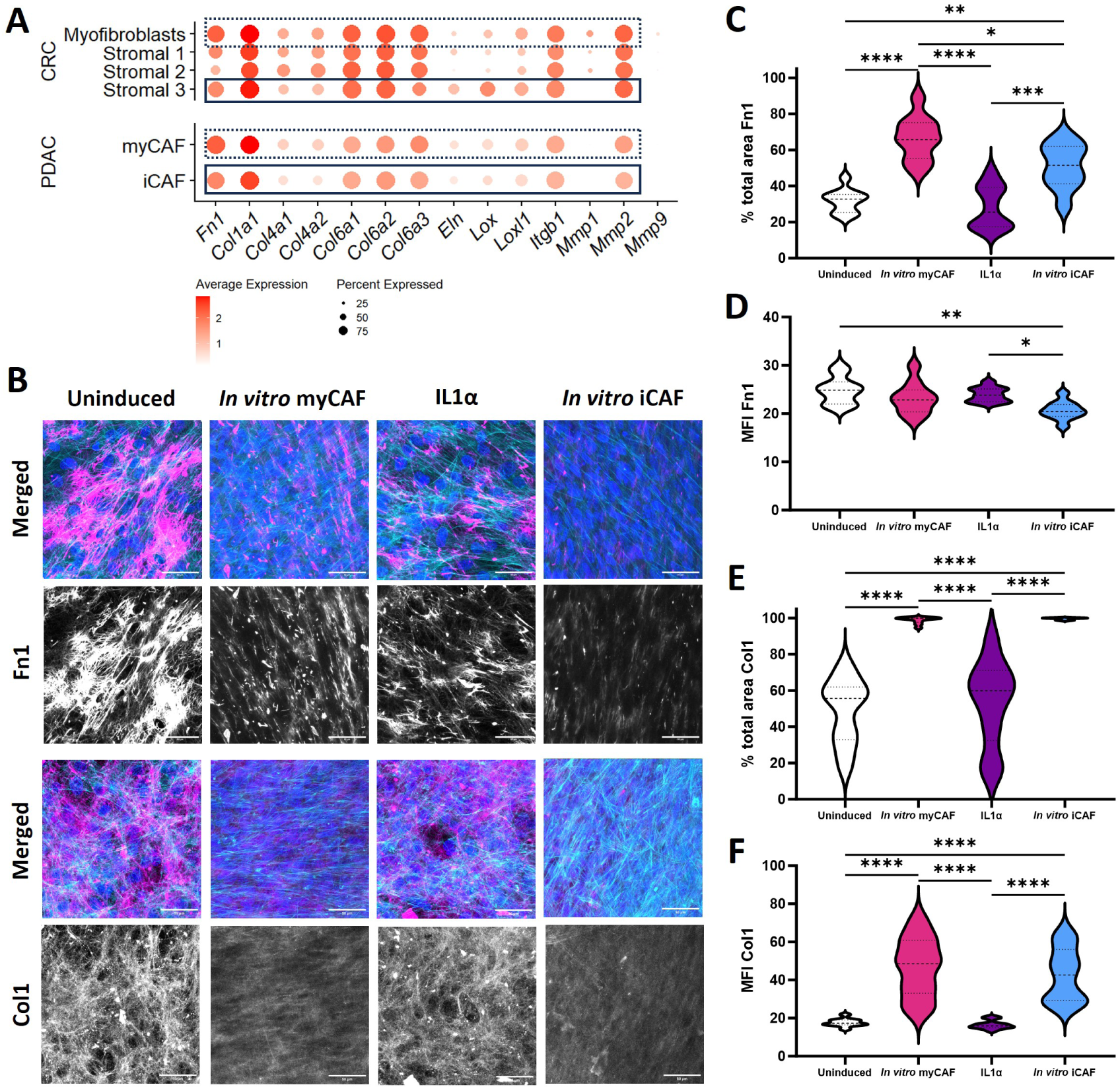
*In vitro* myCAFs and iCAFs produce ECM of comparable composition. **A)** Dot-plots representing average gene expression, and percentage of tumour tissue CAFs expressing *Fn1, Col1a1, Col4a1, Col4a2, Col6a1, Col6a2, Col6a3, Eln, Lox, Loxl1, Itgb1, Mmp1, Mmp2* and *Mmp9*. From the CRC dataset, a comparison is made between subtypes Myofibroblasts, Stromal 1, Stromal 2, and Stromal 3 of tumor origin. From the PDAC dataset, myofibroblastic associated (myCAF) and inflammatory (iCAF) CAFs are compared. **B)** Confocal images of cellularized fibroblast-derived layered matrices fixed and stained with Dapi (blue), phalloidin (cyan) and antibody against Fn1 or Col1 (magenta). Scale bar = 50um. **C-F)** Quantification of % total area covered (C, E) and average mean fluorescence intensity (D, F) of Fn1 and ColI. Data were analyzed by one-way ANOVA followed by Tukey’s multiple comparisons test for multiple images from two independent experiments. *p<0.05; **p<0.01; ***p<0.001; ****p<0.0001.

We next investigated the ECM composition of the *in vitro* CAF subtype-specific CDMs. For that, 12 day matrices were stained for F-actin and nuclei, to identify the cells, and for the ECM components fibronectin (Fn1) and collagen I (Col1) (**Fig. 4B**). Using the fluorescence signal of Fn1 and Col1, we measured the percentage of total area covered by these two ECM components as well as their mean intensity for all the four different CDMs (**Fig. 4C-F**). The *in vitro* myCAF CDMs presented a significantly higher percentage of area covered by Fn1 than the *in vitro* iCAF matrices, followed by significantly lower and comparable area coverage in both uninduced and IL1α-only induced CDMs (**Fig. 4C**). Interestingly, and similarly to the scRNAseq data of tumour-associated iCAFs (**Fig. 4A**), the *in vitro* iCAF CDMs had a generally lower amount of Fn1 than the other conditions, which presented comparable levels of Fn1 (**Fig. 4D**). Col1, however, presented significantly higher area coverage and abundance in both *in vitro* subtype CDMs than in uninduced and IL1α-only induced CDMs (**Fig. 4E,F**). In agreement with the microscopy images and with the scRNAseq data, qPCR confirms that the expression levels of *Col1a1* are equally high in *in vitro* myCAFs and iCAFs both at the beginning of the culture (4 days) and after 2 weeks, albeit over time the levels become lower and more similar to uninduced fibroblasts (**Suppl. Fig. 1B**). Collectively, these observations indicate that, despite some differences, both *in vitro* iCAFs and tumour-resident iCAFs are capable of producing ECM as much as myCAFs do.

We next investigated whether differences in ECM surface coverage and composition would translate into matrices with different biophysical properties, as cell orientation is known to strongly correlate with matrix orientation (52). For this, we fitted the distributions of the orientation of Fn1 and Col1 features with a circular normal distribution (von Mises distribution), providing a mean orientation or peak position and a concentration parameter *k*. In this analysis, a high *k* value indicates highly coaligned structures. The orientation analysis shows that in both uninduced and IL1α-only induced CDMs, Fn1 appears randomly distributed as indicated by the low *k* values (**Fig. 5A**). Conversely, *in vitro* myCAF and iCAF produced a matrix with highly coaligned Fn1, displaying *k* values that are significantly higher (**Fig. 5A**). Also for Col1, we observed that *in vitro* myCAFs generated a collagen matrix that was significantly more aligned than that produced by uninduced or IL1α-only induced cultures. Although the average *k* value of Col1 generated by *in vitro* iCAFs is not significantly different from the other conditions, this value still lies in between the one of myCAFs and uninduced samples (**Fig. 5A**).

**Fig. 5.**
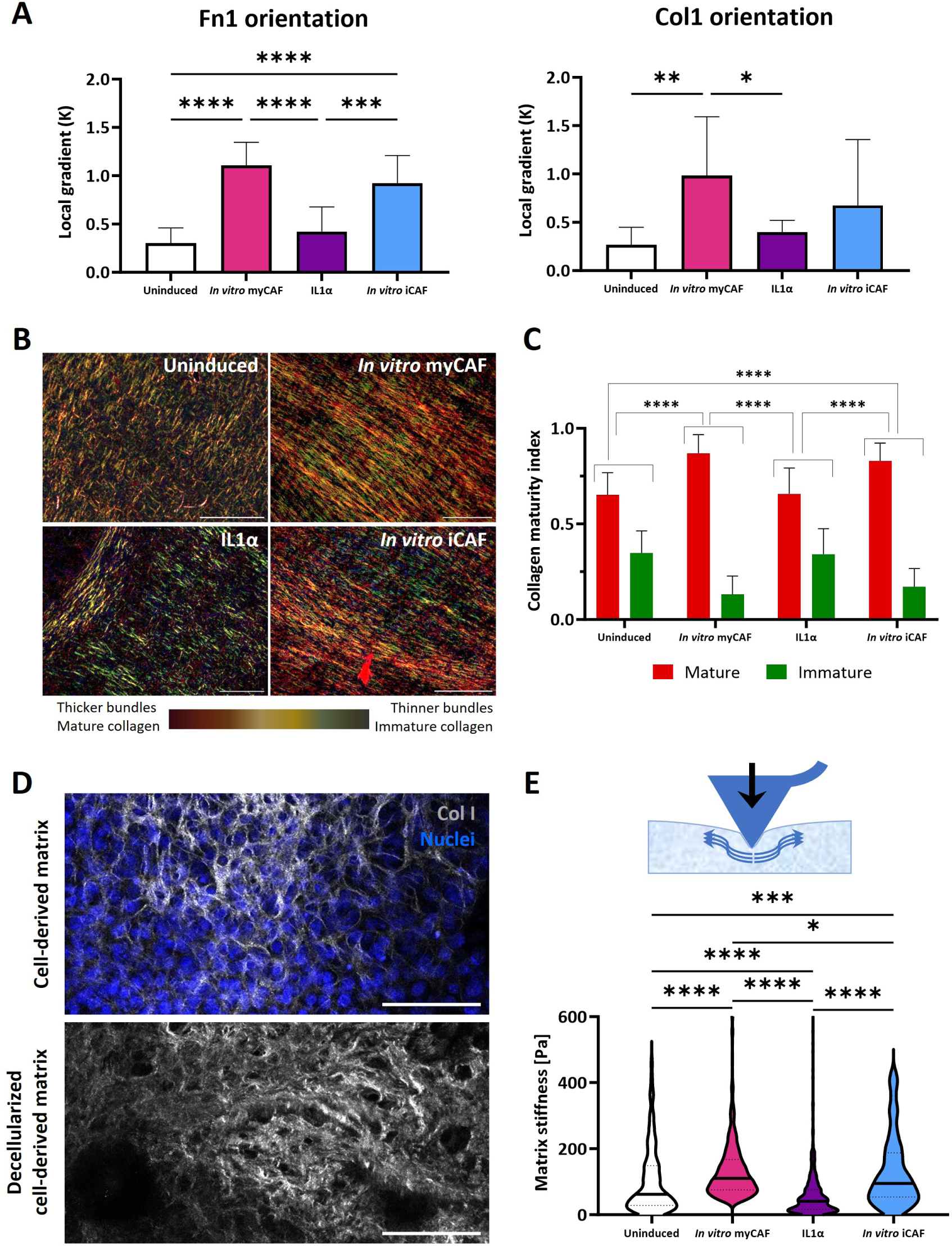
Biophysical properties of fibroblast subtype-derived matrices. **A)** Quantification of matrix directionality for FN1 (left) and COL1 (right) was performed by calculating the Local Gradient (K), which provides the relative directionality of the analysed image, using confocal images of cell-derived matrices after fixation and staining with specific antibodies. Data from two independent samples were analysed with One-way ANOVA followed by Tukey’s multiple comparisons test. *p<0.05; **p<0.01; ***p<0.001; ****p<0.0001. **B)** Picrosirius-red staining of matrices imaged by linearized polarized light. Scale bar = 100um. **C)** Comparison of collagen maturity of different matrices. Measurements represent a quantification of picrosirius red colour and were performed on several images from three independent samples for uninduced, *in vitro* myCAF and IL1α, and two independent samples for *in vitro* iCAF. Data were analysed by a Krustal-Wallis test, followed by Dunn’s multiple comparisons test. ; ****p<0.0001. **D)** Representative images of cellularized (top) and decellularized (bottom) matrix from an uninduced sample obtained by 2-photon fluorescence microscopy. Nuclei were stained with Dapi (blue) while SHG indicates the collagen (gray). Scale bar = 100um. **E)** Cartoon showing how an AFM tip can be used to probe matrix stiffness (top). Multiple measurements (188-542) were performed on several decellularized matrices prepared from three independent samples for uninduced matrices, 4 independent samples for *in vitro* myCAF and IL1α matrices, and one sample for *in vitro* iCAF matrices. A Krustal-Wallis test was performed, followed by Dunn’s multiple comparisons test. *p<0.05; ***p<0.001; ****p<0.0001.

In tissues, collagen fibres are usually characterized by various degrees of supramolecular assembly and maturation. Differently assembled and matured collagens affect tissue microarchitecture and stiffness, hence providing a different mechanical feedback to cells (53–55). To quantify the fractions of mature/immature collagen fibres in the different tissues, we stained the matrices with picrosirius red, and then we observed them under linearly polarized light. This technique enables the visualization of mature collagen bundles in the form of dark orange-red structures, whereas immature fibres are green-yellow (56). Uninduced and IL1α-only induced CDMs were characterized by a large amount of greenyellow signal, whereas *in vitro* myCAFs and iCAFs CDMs contain significantly more mature collagen bundles characterized by the dark orange-red signal (**Fig 5B,C**).

To verify whether such different assembly and maturation of collagen result in different mechanical stiffness of the matrix, we performed force spectroscopy measurements on decellularized CDMs by AFM (**Fig. 5D,E**). The decellularization protocol we adopted liberated the CDMs from the cellular component with a negligible influence on the collagen network, as visualized by SHG microscopy (**Fig. 5D**). Decellularization enabled us to measure the actual stiffness of the matricellular components of the *in vitro* generated tissue. In agreement with the collagen microarchitecture and maturation state, the most compliant tissues were generated by the uninduced and IL1α-only induced fibroblasts, while the stiffest tissues were produced when TGFβ was supplemented in the culture (i.e. myCAFs and iCAFs) (**Fig. 5E**).

Collectively, these results demonstrate that CAF subtype-specific CDMs can be generated *in vitro* and that our *in vitro* iCAFs and myCAFs are equally capable of generating thicker matrices with well-defined alignment and higher stiffness that are reminiscent of tumour stromal ECM.

### Fibroblast subtype-specific CDMs induce upregulation of Arg1 in co-cultured macrophages

Within the TME, CAFs and their ECM are known to influence other tumour-resident cell types such as the heterogeneous population of tumour-associated macrophages (TAMs) (57, 58). Even though recent studies delved into the effect of the tumour stroma ECM on TAM polarization (59), the different effects of myCAF- and iCAF-generated matrices on macrophages are yet to be thoroughly investigated. We therefore sought to determine whether our *in vitro* CAF subtype-specific CDMs were capable of influencing macrophage phenotype. For this, we generated the CDMs under the four different conditions, washed with plain medium, seeded unstimulated bone-marrow derived macrophages (BMDMs) (60) on top and after 2 days of coculture we used flow cytometry to measure the expression of Arginase-1 (Arg1), which is known to be upregulated in pro-tumorigenic macrophages (61, 62) (**Fig. 6A**). Specifically, we cocultured the BMDMs with either untreated (i.e. live fibroblasts) or mitomycin C (MMC)-treated (i.e. non-proliferative fibroblasts) (63) CDMs to determine whether the presence of active fibroblasts within the CDMs is needed to influence the BMDM phenotype. As shown in **Fig. 6B-C**, Arg1 expression is higher in BMDMs seeded on *in vitro* myCAF and *in vitro* iCAF CDMs, with respect to the BMDMs seeded on CDMs of uninduced or IL1α-only induced fibroblasts, regardless the physiological state of the fibroblasts (live or MMC-treated). This is shown not only by the values of mean fluorescence of the entire BMDM populations but also by the percentage of BMDMs expressing high levels of Arg1.

**Fig. 6.**
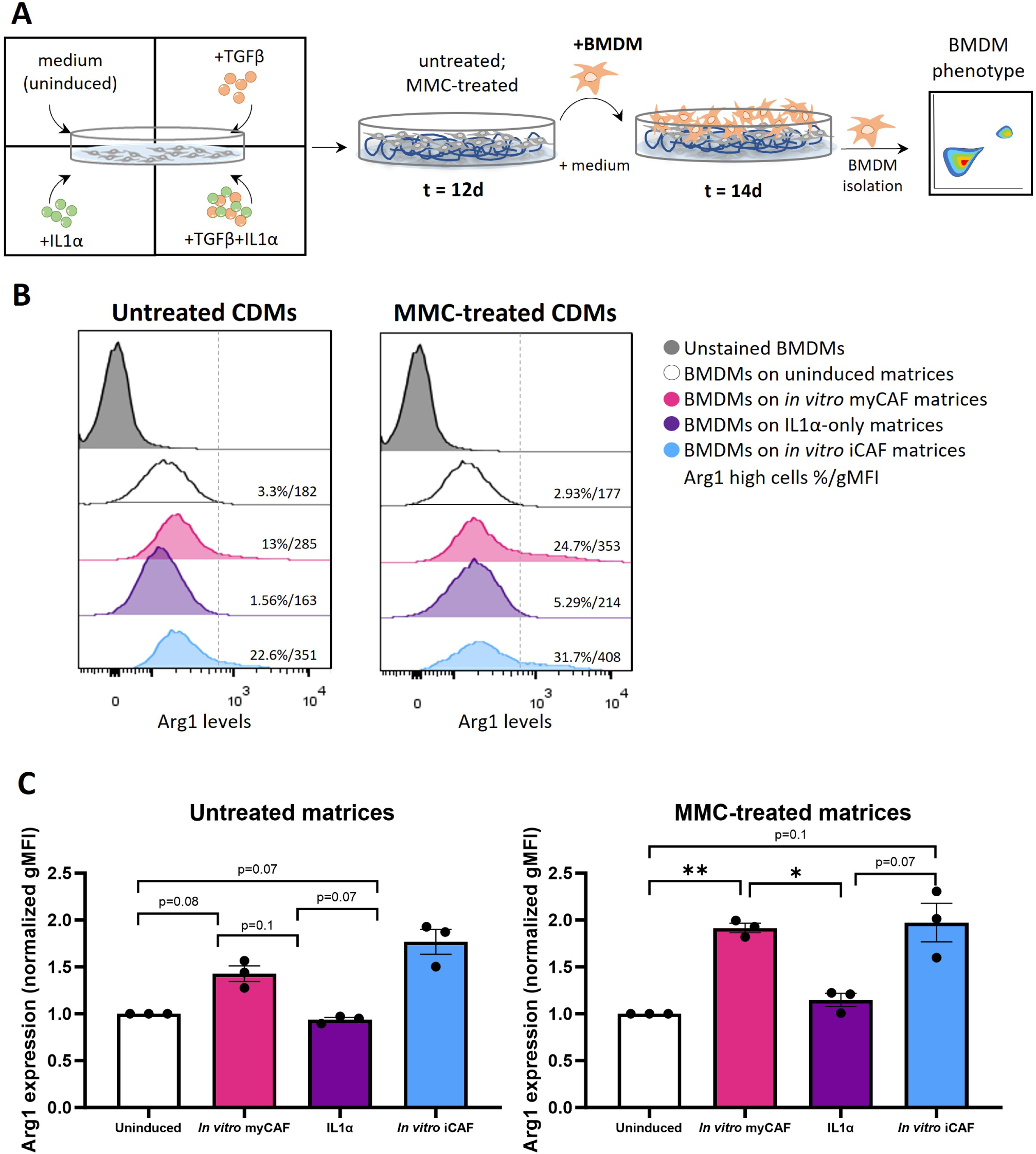
*In vitro* fibroblast subtype-derived cellularized matrices enhance Arg1 expression in co-cultured macrophages. **A)** Schematic representation of the experimental conditions. After 12 days of culture, the cellularized layered matrices were washed with medium and either left untreated or treated with MMC to inhibit fibroblast duplication and secretion. BMDMs were added to the different matrices, co-cultured for 2 days and harvested to determine their phenotype. **B)** Histograms showing the expression levels of Arg1 on BMDMs that were cocultured for 2 days on different matrices. Respresentative repeat of NS3. **C)** Geometric mean of Arg1 expression of BMDMs cultured for 2 days on different matrices. Data from 3 independent samples, analyzed with One-way ANOVA followed by Tukey’s multiple comparison test. *p<0.05; **p<0.01

These results suggest that these fibroblast subtypematrices specifically provide signals that lead to increased Arg1 levels on the BMDMs and indicate that these fibroblast-subtype specific CDMs, either live or MMC-treated, are amenable as *in vitro* stromal tissues for future mechanistic studies to unravel pathways regulating fibroblast-immune cell crosstalk.

## Discussion

In this work, we generated a 3D *in vitro* model of fibroblast subtypes that resemble tissue-remodelling myofibroblast-like myCAFs and inflammatory fibroblast-like iCAFs. We used these *in vitro* subtypes to characterize their role on ECM production, deposition and remodelling. We found that TGFβ is needed to induce a myCAF-like phenotype on NIH-3T3 fibroblasts, while the iCAF-like phenotype can be achieved by a combination of TGFβ and IL1α and that these in vitro fibroblast subtypes are phenotypically very similar to the CAF subtypes characterized in tumour sections of cancer patients. Moreover, these two *in vitro* fibroblast subtypes are equally able to generate layered and stiff matrices with aligned and mature ECM, demonstrating for the first time that inflammatory fibroblasts may also exhibit ECM-remodelling capacity. Finally, matrices generated from these *in vitro* fibroblast subtypes, but not from uninduced or IL1α-only stimulated fibroblasts, enhance expression of the immune suppression marker Arg1 in co-cultured BMDMs and are therefore suitable for mechanistic studies about cell-cell interactions between fibroblast subtypes and other stromal components.

Correlating the expression of established myCAF and iCAF markers in our *in vitro* subtypes with the analysis of publicly available scRNAseq data of CRCand PDACassociated CAFs (36, 37), we conclude that *in vitro* myCAFs can be induced by TGFβ alone, while a combination of TGFβ and IL1α is necessary to mimic the *in vivo* iCAF phenotype. Although upregulation of *Acta2* and *Tagln* is widely used as a myCAF signature, the scRNAseq data clearly indicates that almost all fibroblasts populating either the tumour or the surrounding normal tissue have similar expression levels of these two genes. On the contrary, higher levels of *Ccl2* and *Il6* in combination with the specific *Lif* expression unequivocally distinguish iCAFs from myCAFs. In our *in vitro* subtypes, *Ccl2, Il6* and *Lif* can be induced by IL1α alone, but only when IL1α and TGFβ are added together to the fibroblasts we observe further increase of these markers as well as induction of *Acta2* and *Tagln*, as found in *in vivo* iCAFs from CRC and PDAC. The presence of *Acta2* and *Tagln* in both myCAFs and iCAFs nicely correlates with the observation that the *in vitro* subtypes are equally capable of contracting fibrillar collagen matrices. It is however important to point out that we have benchmarked our *in vitro* fibroblast subtypes to only two publicly available databases of scRNAseq of tumour tissues. As several gene expression datasets exist for the same cancer type (28, 29) and different datasets are available for cancer types other than CRC and PDAC (26, 27), we find opportune to emphasize that some heterogeneity in the expression levels of subtype-specific markers is to be expected between CAF subtypes from different tumours but also between different datasets of the same tumour type. For example, myCAFs from PDAC present high levels of *Thy1* compared to iCAFs and apCAFs (29), while in breast cancer, all fibroblast subtypes present similar levels (26, 46). On the other hand, even though iCAFs are reported not to express any *Tagln* in breast cancer (26), iCAFs in PDAC have been shown to express *Tagln*, at the same or slightly lower level as compared to myCAFs (29, 37). Another example is IL6, classically considered an iCAF marker, which is also expressed in myCAFs across three cancer types (i.e. melanoma, head and neck squamous cell carcinoma and lung cancer) (27). Moreover, tissue-remodelling myofibroblasts found in cancer lesions may slightly differ from the same subset found in fibrosis or inflammation (1, 3). Our *in vitro* subtype induction model based on TGFβ and IL1α cannot possibly recapitulate this heterogeneity but does offer a robust and easily adaptable system to try combinations of other inflammatory cytokines and growth factors known to be secreted in the TME or in other pathological tissues.

Our current findings expand previous studies, which support a mutually exclusive polarization hypothesis by TGFβ and IL1α (6, 31, 32). Biffi and colleagues (32) show that IL1α inhibition boosts a pro-myofibroblastic phenotype in pancreatic stellate cells (PSCs). Furthermore, they report an inhibitory effect of TGFβ on IL1α-preconditioned PSCs, which resulted in lower iCAF polarization (32). Indeed, in our subtype induction model we observed that in fibroblasts stimulated with IL1α only, expression of typical myCAF markers decreased. However, the simultaneous stimulation with TGFβ and IL1α seems to have a synergistic pro-inflammatory effect on fibroblasts, which is in contrast with this previous finding but could be explained by differences between the PSCs used by Biffi and colleagues and the NIH3T3 fibroblasts used in our study. Further investigation is needed to better elucidate how the interplay between the TGFβ and the IL1α pathways impacts phenotype plasticity of fibroblast subtypes. In those previous studies, tumour CM, which contains miscellaneous secreted cytokines and other biochemical signals, was also used to stimulate PSCs resulting in induction of iCAFs (6, 32). However, the outcome of this stimulation can be quite diverse as different tumours produce different CMs, which are usually not thoroughly characterized, resulting in variable potency when inducing CAFs. We here propose a standardized and robust model of fibroblast subtypes induction based on TGFβ and IL1α that can reproducibly induce *in vitro* tissue-remodelling fibroblasts and inflammatory fibroblasts, both in 2D and 3D culture conditions, that highly resemble their *in vivo* CAF counterparts.

Analysis of the scRNAseq databases also revealed that *in vivo* iCAFs express genes related to ECM production, such as collagens and fibronectin, and to ECM remodelling and cross-linking, such as metalloproteinases and lyxyl oxidases respectively, at a comparable level as in myCAFs. Similarly, the TGFβ and IL1α combination promotes expression of Col1 at comparable level to TGFβ alone. This observation correlates well with our finding that both fibroblast subtypes generate thick, aligned matrices likely due to the contractile forces they exert on the ECM fibres. Fn1 and Col1 are two well-characterized ECM components, associated with early and late desmoplastic stages respectively (64, 65). Indeed, they are known to have a sequential deposition, as Fn1 acts as a scaffold for Col1 deposition (49, 66). Interestingly, we did observe slightly less Fn1 in the matrices of inflammatory fibroblasts. Related to this observation, Qi and colleagues showed that IL1α secreted by macrophages correlated with lower expression of Fn1 in FAP+ fibroblasts in colorectal cancer (67). Despite presenting comparable Col1 coverage and maturity to tissue-remodelling fibroblast matrices, inflammatory fibroblast matrices exhibited a minor decrease on Col1 intensity and were slightly softer. As Fn1 serves as a scaffold for collagen deposition and assembly (49), less Fn1 may allow for a smaller fraction of collagen to be assembled, which may result in softer matrices. Indeed, a recent collagen profiling study of myCAFs and iCAFs throughout different cancer types showed that myCAFs express slightly higher levels of Col1 than iCAFs, which could be explained by IL1α inhibitory effects on pro-fibrotic TGFβ induction (33, 41, 43, 44). These findings are in line with the different composition and stiffnesses of our *in vitro* subtype-specific matrices.

Fibroblasts and their ECM are known to influence the behaviour of neighbouring cells, in particular macrophages, in health and disease (68). CAFs have been extensively described to be in close contact with TAMs, both cell types synergizing within the stromal compartment to generate a protumorigenic environment (40, 57, 69, 70). Interestingly, we found that Arg1, a classical pro-tumorigenic, immune suppressive TAM marker (71), is clearly upregulated on BMDMs when cultured on both live and MMC-treated matrices and not observed when BMDMs are cultured on matrices of uninduced or IL1α-only stimulated fibroblasts. Classically, IL4 stimulation significantly increases Arg1 upregulation in TAMs, likely due to the fact that IL4 promotes TGFβ2 expression, which can have an autocrine effect, inducing Arg1 expression and, at the same time, TGFβ1 promotes expression of IL4 receptor (72–75). On another note, increased stiffness can also promote Arg1 expression (69). Our findings that even in the absence of IL4, BMDMs upregulated Arg1 specifically on the matrices of the two fibroblast subtypes are supported by recent discoveries where TGFβ and stiffness are described to synergize and promote Arg1 expression on BMDMs in breast cancer (59). We hypothesize that there might be available TGFβ in both subtype-derived matrices, either secreted or ECM-anchored, which induces BMDMs to upregulate Arg1 production. Of course more detailed studies are needed to depict the potential of these fibroblast subtype specific matrices in inducing pro-tumorigenic features in cocultured immune cells. What we show is a useful and versatile heterocellular coculture system that could be adapted to test more in detail, next to Arg1, possible changes of other immune suppressive markers in macrophages, or other immune cells, cocultured on these fibroblast subtype matrices. Finally, as macrophages can also remodel the ECM, the effects of differently polarised macrophages on the fibroblast subtypes trapped in their own matrices could also be investigated.

In the context of therapeutic design and optimization, these subtype specific matrices and the various cell types involved could also be used for drug screening purposes. Arg1, a classical pro-tumorigenic TAM marker, catabolizes L-arginine, an essential amino-acid, into L-ornithine, subsequently promoting amino-acid starvation in the TME, which restricts T cell activation and proliferation (71).

Targeting specific fibroblast subtypes may offer new treatment options in several pathologies. CAFs have been in the spotlight in recent years regarding the uncovering of their relevance and role in cancer evolution, especially gaining importance as the main key player of the TME. Several approaches have been developed to hinder their functions and promote an anti-tumoral environment. CAF-specific targeting efforts have been focused either on targeting the CAFs themselves, or the ECM (76–78). Specific myCAF depletion experiments, however, resulted into a faster tumour development and lower patient survival (79). This can be explained by the myCAF characteristic behaviour of building up a capsule around tumour clusters, which has an organic anti- and pro-tumorigenic effect by either impeding tumour cell access to immune cells or chemotherapeutic molecules, and simultaneously restraining tumour growth and migration (80). Moreover, based on our data, iCAFs remaining after myCAF depletion are well able to deposit and remodel the ECM. Targeting the differential ECM, thus, might present successful results, and that is why it is important to understand differences between subtypes. However, those differences portrayed here, such as differential Fn1 composition, might be too minor to serve as a CAF-subtype specific ECM target per se. In any case, more in-depth studies of additional interesting components of the ECM, such as other structural proteins or molecules anchored to the matrix, might shed light into further differences between CAF subtype matrices, serving as a platform for identification of new targets.

Inhibiting CAF activation or re-polarization of CAFs into healthy fibroblasts might be the better compromise, and experimental models are already yielding promising results (81, 82). As the biochemical composition of the TME is highly complex and likely changes over time, foci of specific cytokine secretion by the tumour or the immune cells or by the CAFs themselves could locally induce changes in CAF subtype phenotypes: a release of IL1α by TAMs in a TGFβ rich TME area could induce the local myCAFs to become iCAFs. Detailed experiments of fibroblast lineage tracing within the TME or in fibrotic lesions aimed at unravelling CAF plasticity are warranted. Similar mechanisms may be occurring in fibrotic or inflamed tissues, where the tissue-remodelling and the inflammatory phenotypes may not be mutually exclusive but be dictated by the local levels of inflammatory factors of inflammatory cells. The availability of *in vitro* models of fibroblast subtypes like the one proposed here will contribute to a better understanding of this heterogeneous population and identify potential new targets for therapeutic development, besides constituting a platform to model other emerging fibroblast subtypes and investigate their influence on neighbouring cells. Finally, besides altering the biophysical properties of the tissue, pathological ECM plays other functions that we are only beginning to understand. For example, ECM fragments released by proteolytic enzymes can bind integrins from cancer cells thereby reducing their proliferation and invasive capacity (24). Also, ECM uptake and catabolism can be a source of nutrients for cancer cells under amino-acid starvation conditions (83). Whether different fibroblast subtypes play different roles in these processes and whether these ECM functions are also important in other pathologies besides cancer are still open questions. The *in vitro* fibroblast subtypes presented in this study offer a robust and tractable model to investigate these mechanisms.

## Experimental Procedures

### Analysis of single-cell RNA sequencing data

#### Data used in this study

We used scRNA-seq and metadata from the CRC (36) and PDAC (37) datasets. The read counts and associated metadata of the CRC dataset were retrieved from the NCBI Gene Expression Omnibus (GEO), accession code GSE132465, containing 63,689 cells from 23 colorectal cancer (CRC) patients with 23 primary colorectal cancer and 10 matched normal mucosa samples. We used 5,933 cells from the CRC dataset that were annotated by the authors as “stromal cells”. The PDAC data were retrieved from the Zenodo repository (3969339). They contain 57,530 cells from 3 control pancreases of non-pancreatic cancer patients and from 8 pancreases from patients with a non-malignant pancreatic tumour. From these 57,530 cells, we used a total of 6,926 cells that were labelled by the authors as fibroblasts.

#### scRNA-seq analysis

The Seurat package (84) was used to analyse the two datasets in R (R core team). The CRC data were first normalized using *NormalizeData*, and the 2000 most variably expressed genes were determined with *FindVariableFeatures*. The data was scaled by carrying out a linear transformation using *ScaleData*. Based on the 2000 variably expressed genes, a principal component analysis was run using *RunPCA*, to compute the first 15 PCs used in the downstream analysis. Euclidean-based distances were calculated between the cells and were used to find the k-nearest (k=20) neighbours from the PCA space using *FindNeighbors*. The Louvain clustering algorithm was used to the group cells together into clusters using *FindClusters*. A TSNE plot was generated to visualize the clustered data in a low dimensional space, using *RunTSNE*.

Upon closer inspection of the CRC cells that were annotated as “stromal” cells we noticed that it also contained smooth muscle cells and Tip-like ECs, which we excluded as the only cells relevant to the study are the CAFs/stromal cells. We then selected clusters that contained sufficient numbers of CAFs/Stromal cells of both tumour and non-tumour origin for comparison, totalling to 3,392 cells. The data was then re-clustered using only these cells by repeating the above processing steps, and split based on “tumour” or “normal” classification in the metadata to generate the dot plots with the expression levels. With respect to the PDAC data, as they were only partially annotated, the single cell atlas of the PDAC tumour microenvironment (45) was used for deeper annotation. Specifically, the “Stroma_Subset2021” reference data (github) was used to identify relevant tumour and non-tumour associated fibroblasts. The fibroblasts were annotated by first using *FindTransferAnchors* that identifies cells of similar gene expression profiles (anchors) between reference and query single cell data. These anchors were then transferred to the query PDAC data using the function *TransferData* to classify the PDAC cells based on the reference data. The inferred cell types were then added to the PDAC dataset as metadata for further analysis of CAFs and stem cells. The annotations were examined for the expression of key marker genes of each cell type to confirm the cell type annotations. Using the generated annotations from the PDAC annotation, we included only complement secreting CAFs (csCAFs), iCAFs, myCAFs, quiescent pancreatic stellate cells (qPSCs), and smooth muscle pancreatic stellate cells (smPSCs). These cells were then normalized, variable features were computed, the data was scaled, a PCA was run, neighbours and clusters were found as described above. Using the metadata, the cells were then split into normal and tumour conditions for the comparisons.

#### Visualization

The Seurat package was used to generate the TSNE plots, which were combined using the patchwork package. Genes of interest LIF, CCL2, IL6, TAGLN and ACTA2 were selected for CAF subtype classification, while MMP9, MMP2, MMP1, ITGB1, LOXL1, LOX, ELN, COL6A3, COL6A2, COL4A2, COL4A1, COL1A1 and FN1 were selected as ECM-related genes of interest.

#### Software

For data processing and visualization R (v 4.3.2), Seurat (v 5.0.2), dplyr (v 1.1.4), cowplot (v 1.1.3), patchwork (1.2.0) were used.

#### Data availability

The code used for the analysis and visualization of the data is available on GitHub (https://gitlab.cmbi.umcn.nl/cmbi/cafs_sc_analysis).

### Fibroblast culture conditions

Embryonic mouse fibroblast cell line NIH3T3 was obtained from ATCC (CRL-1658) and cultured in high-glucose DMEM with 4.5 g/L glucose (Gibco, #10938025) supplemented with 10% fetal calf serum (FCS) (Cytiva, SH30541.03), penicillin-streptomycin (Sigma-Aldrich, #P4333), 200 mM L-glutamine (Gibco, #25030-024) and 1% sodium pyruvate (Gibco, #25030-024), namely complete medium. Cells were washed 1X PBS (phosphate buffered saline, in-house production), collected using TrypLE Express Enzyme (Gibco, #12604-013) and centrifuged 5 minutes at 250G and further resuspended in complete medium. Cells were passaged 1:10. Cells were incubated at 37°C with 5% CO2 between experiments or passaging.

### Differentiation of fibroblasts in 3D fibrillar collagen gel

Cells were collected and resuspended in 1.7 mg/mL of neutralized Rat Tail Collagen Type I (Corning® Life Sciences, #354249). Drops were polymerized at 37°C for 30 minutes upside down to ensure a 3D structure. Induction medium was added to the drops, which consists of 5% FCS, penicillin-streptomycin, L-glutamine and 1% sodium pyruvate in high-glucose DMEM, further complemented by 5 ng/mL of TGFβ (HEK293 derived) (PeproTech, #100-21-10UG) and/or 1 ng/mL of murine IL-1α (PeproTech, #211-11A-10UG).

### RNA isolation and RT-qPCR

Collagen cell-containing drops were collected, washed with PBS and dissociated using Collagenase I (Sigma, C0130-1G) for 10 minutes at 37°C. The same pipeline was performed when collecting RNA from matrices. Cells were further lysed using the lysis buffer of Aurum™ Total RNA Mini Kit (Bio-Rad, #7326820). Lysates were passed 10 times through a sterile BD Microlance™ 3 needle (#300635, 27G 0.4 × 13 mm) attached to a 1 mL BD Luer-Lok^™^ Syringe (#309628) and RNA was further isolated using the protocol provided by the manufacturer. For cDNA, the iScript™ cDNA Synthesis Kit (Bio-Rad, #1708891) was used. RT-qPCR was executed with iQ™ SYBR Green Supermix (Bio-Rad, #1708886) on the CFX96™ Touch Real-Time System. Primers (Table1) were used for gene expression of markers, and normalized to the housekeeping genes *Ppia* and *B2m*.

Table 1: Primer list for CAF subtype markers Forward (5’-> 3’) Reverse (5’-> 3’) Ppia CAAACA-CAAACGGTTCCCAG TTCACCTTCCCAAAGAC-CAC B2m GGTCTTTCTGGTGCTTGTCT ACG-TAGCAGTTCAGTATGTTCG Acta2 GGACGTACAACTG-GTATTGTGC TCGGCAGTAGTCACGAAGGA Tagln GTGTGATTCTGAGCAAATTGGTG ACTGCTGC-CATATCCTTACCTT Col1a1 GCTCCTCTTAGGGGC-CACT CCACGTCTCACCATTGGGG Il6 CTGCAA-GAGACTTCCATCCAG AGTGGTATAGACAGGTCT-GTTGG Ccl2 TTAAAAACCTGGATCGGAACCAA GCATTAGCTTCAGATTTACGGGT Lif AACCAGAT-CAAGAATCAACTGGC TGTTAGGCGCACATAGCTTTT

### Collagen ring contraction assay

Wells of 24 well-plate (VWR Tissue Culture Plates, 10062-896) were coated with sterile 0.1% BSA/PBS (Sigma-Aldrich, A9647) to allow for collagen to detach once the medium was added. Cells seeded at a density of 5×10^4^ cells/mL in 1.7 mg/mL of neutralized rat tail collagen. 120 μl of the cell mixture was added on the edges of the well. After polymerization, induction medium was added, supplemented with 5 ng/mL of TGFβ and/or 1 ng/mL of IL1α or 10 μM Rock Inhibitor (Selleckhem, S1049, Y-27632). After 2 days, rings were fixed in 4% PFA (Merck, 16005) for 30 minutes, washed 3x with PBS, mounted on a slide and images were taken using a smartphone camera (Samsung Galaxy A71). The collagen area of each ring was quantified using FIJI (version 1.8.0).

### CAF subtype-specific matrix production

Cells were collected using TrypLE and plated on top of either an EtOH-sterilized coverslip or a layer of polydimethylsiloxane (PDMS, Sylgard 184, Dow Corning, Midland, MI) at a concentration of 8×10^4^ cells/mL. Complete medium was used, supplemented with 100 μg/mL Ascorbic Acid (2-O-α-D-Glucopyranosyl-L-ascorbic Acid, Tokyo Chemical Industry, #G0394) and 5 ng/mL of TGFβ and/or 1 ng/mL of IL1α. To test whether cells within matrix-producing cultures were still conserving subtype-specific gene expression, cultures were kept for 4 and 14 days, isolating RNA and performing RT-qPCR as previously described.

### Mechanical characterization of decellularized cell-derived matrices

At the end point of the culture, cell-derived matrices (CDMs) were decellularized prior to Atomic Force Microscopy (AFM) characterization. CDM decellularization was performed by incubating the samples for one hour in a solution containing: Triton 1% (Sigma), EDTA 20 mM (Sigma) and NonidetP40 Substitute (Fluka) 1% in PBS (Sigma). Samples were then thoroughly rinsed with PBS. The nuclei were previously stained with Hoechst (33342, Invitrogen 1:10000) 15 min at 37°C in order to visually assess the effectiveness of the decellularization process. The decellularized CDMs (dCDMs) were then stored in PBS at 4°C. The mechanical properties of the dCDMs were measured using a JPK NanoWizard II AFM, using a silicon nitride cantilever (SAA-SPH-5UM, Bruker) with a spherical tip (radius 5 μm). The cantilever spring constant was estimated using the thermal fluctuation method. Cantilever spring constant (k) was measured experimentally and values generally fell within the 0.08-0.10 N/m range (nominal spring constant 0.08 N/m as per manufacturer’s indication). Force curves were recorded at different points of the matrices on regions of interest of 100 × 100 μm area, at a scanning frequency of 1 Hz and a velocity of 2 μm/s. Over 60 force curves were measured on each area, at room temperature in PBS. The mechanical properties, in terms of Young’s modulus (E), were estimated by fitting the force-indentation curve to the Hertzian model for linearly elastic materials and spherical indenter using JPK Data Processing software (85). The matrices were assumed to be nearly incompressible with a Poisson’s ratio of 0.5. The Young’s modulus was found for each force curve using the fitting algorithm provided by the JPK software. A maximum indentation of approximately 400 nm was chosen for fitting. Average Young’s modulus was computed by averaging all force curves for a given condition.

### Picro-Sirius Red staining

To visualize collagen fibres, fixed CDMs were immersed in a Picro-Sirius Red solution, made of 0.1% w/v Sirius Red (Direct Red 80, Sigma-Aldrich) in saturated aqueous solution of picric acid (1.3% in water, Sigma-Aldrich) for 45 minutes at room temperature and then rinsed twice in a 1% acetic acid solution in water. Stained CDMs were visualized under linearly polarized light with a 10X objective mounted in an upright microscope (Olympus, BX53). In order to quantify the coloured sections with DirectRed80, the captured images were then analysed with the Fiji software (version 1.54f) using the ‘CMM_PR-BRF_Analyser’ macro (86). This script allowed us to automate the quantification of the birefringent signal area in the “red-orange” (colour scale 0-27, 230-255) and “yellowgreen” (colour scale 28-140) channels of each image that roughly correspond to mature collagen fibres and immature collagen fibres, respectively.

### Orientation analysis

Images of cells and collagen in CDMs and dCDMs were collected fluorescence microscopy and analysed by ImageJ software. To assess the orientation of collagen fibres and the arrangement of cells within the matrices, the acquired images were analysed with the Fiji software using the “Directionality” plugin. To describe the fibre distribution as a function of the fibre angle we used a modified π-periodic Von Mises distribution(87). The fibre density function *ρ*(*ϑ*) is written as

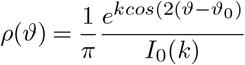

in which *k* describes the concentration of the fibre distribution around the mean angle *ϑ*_0_, *I*_*0*_ is the modified Bessel function of the first kind of order 0. *k* and *ϑ*_0_ were obtained by fitting the Von Mises distribution with the distribution obtained via image analysis. Two images, in the form of a 2×2 tile scan, were acquired per each of the four conditions (n.s. 3, for a total of 6 tile scans per condition).

### Measurement of matrix thickness

To evaluate matrix thickness in each condition, three z-stacks for each of the three independent samples were analysed using the Orthogonal Views function in Fiji software. The thickness was measured with the command “measure” at 3 different locations (left-centre-right) of the z-stack. The fluorescence intensity signal profile was evaluated by first applying a mask generated with Otsu threshold and then measuring the average intensities of the images in the xy plane constituting the z-stack. The thickness of each z-stack was normalized with respect to the maximum height, thus obtaining intensity profiles in the 0-1 range.

### Immunofluorescence staining and image acquisition and quantification of ECM components

CDMs were grown for approximately 12 days, fixed with 4% PFA, washed 3x with PBS, blocked with 20mM Glycin/PBS (MP Biomedicals, 808831) and stained with different combinations of Dapi (Sigma-Aldrich, #D9542), phalloidin-633 (Thermofisher Scientific, A22284), anti-fibronectin antibody (Sigma-Aldrich, #F3648) and anti-collagen I antibody (Cell Signaling Technology, 72026). Z-stacks were taken from all stained CDMs using a 40x water immersion objective on the LSM880 confocal microscope (Zeiss) and images were further analysed using FIJI software. To observe the distribution and organization of collagen fibres, z-stack images of CDMs and dCDMs were collected with a Leica TCS SP 5 (Leica Microsystems) combined with MPM in which the NIR femtosecond laser beam was a tunable compact mode-locked titanium:sapphire laser (Chameleon Compact OPO-Vis, Coherent). Two-photon excited fluorescence was used to induce the second harmonic generation (SHG) signal of collagen. The samples were observed by setting the excitation at 840 nm and the emitted signal was collected in the 415–425 nm range. The SHG images were acquired with a resolution of 12 bit, 512×512 pixels by using a 25× water immersion objective (HCX IRAPO L n.a. 0.95).

### BMDMs and CDMs coculture

Bone marrow was isolated from the tibia and femur of C57BL/6J mice. Bone marrow was cultured for 7 days in non-tissue culture treated dishes in RPMI-40 (Gibco, 42401-018), supplemented 10% FCS, 1% v/v ultraglutamine-1 (Lonza, #BE17-605E/U1), 0.5% v/v 100X Anti-Anti (Gibco, #15240-062)), and 25 ng/mL M-CSF (Peprotech, #300-25) to achieve macrophage differentiation. After 7 days, bone marrow-derived macrophages (BMDMs) were harvested with 0.2 mM EDTA (Invitrogen, #15575020) for 1 h at 4°C. After washing with PBS, BMDMs were seeded with at a density of 200.000 cells on fibroblast-derived matrices. Prior to BMDM seeding, fibroblast-derived matrices were either washed with PBS or treated with 4 μg/mL mitomycin C (MMC) for 2 h at 37C. BMDMs were then cultured on the matrices for 48 h in the presence of 20 ng/mL M-CSF. and then collected from the matrices by using 0.2 mM EDTA at 4°C. After harvesting, cells were washed with PBS and Fc receptors were blocked for 25 minutes with CD16/32 block (1:200, Tonbo Biosciences, #70-0161-M001). Subsequently, cells were stained with anti-F4/80 antibody (BM8, Biolegend, #123110, 1:100) to identify the BMDM population. After 20 minutes, cells were washed and stained for viability using LIVE/DEAD Fixable Yellow (Invitrogen, #L34959). To allow for intracellular Arg1 (A1exF5, Invitrogen, #173697-82, 1:100) staining, cells were permeabilized and fixed with the eBioScience Fix/Perm kit (Invitrogen, #00-5123-43 #00-5223-56) according to the manufacturers protocol. After staining, cells were washed and flow cytometric analysis was performed on the BD FACS Lyric. Data were analysed using FlowJo (version 10).

### Statistical analysis

Visualization of results and statistical analyses were done with the software GraphPad Prism (version 10.1.2). Statistical tests and significance are stated in the figure legends. *p<0.05, **p<0.01, ***p<0.001, ****p<0.0001.

### Data availability

All primary data supporting the conclusions made are available from the authors on request.

## ACKNOWLEDGEMENTS

This work was financially supported by an NWO-XS grant (OCENW.XS23.2.137) awarded to A.C. and an NWO KLEIN grant (OCENW.KLEIN.494) awarded to K.D.. F.J.O. and E.J. were financially supported by the Intramural Radboudumc Hypatia funding. The authors also thank the Radboudumc Technology Center Microscopy for the use of their microscopy facilities.

## AUTHOR CONTRIBUTIONS

F.J.O., C.C., M.V. and A.C. designed the experiments. F.J.O., C.C., M.V., E.J. and V.W. performed the experiments and analysed the data. K.G. and M.A.H analysed the scRNAseq datasets. M.Z. and K.D. provided advice on data analysis. F.J.O., C.C., M.V. and A.C. wrote the manuscript with input from all coauthors. A.C. supervised the project.

## DECLARATION OF GENERATIVE AI AND AI-ASSISTED TECHNOLOGIES IN THE WRITING PROCESS

During the preparation of this work the authors did not used any AI and AI-assisted technologies.

