## Supplementary Information for "*In vitro*-generated inflammatory fibroblasts secrete extracellular matrix with biochemical and biophysical properties similar to tissue-remodelling fibroblasts"

[1]Department of Medical BioSciences, Radboud university medical center, Geert Grooteplein Zuid 26-28, 6525 GA Nijmegen; [2] Department of Chemical, Materials and Industrial Production Engineering, University of Naples Federico II, P.le Tecchio 80, 80125, Naples, Italy, [3]Center for Advanced Biomaterials for Healthcare@CRIB, Fondazione Istituto Italiano di Tecnologia, Largo Barsanti e Matteucci 53, Naples, Italy, and [4]Interdisciplinary Research Centre on Biomaterials , University of Naples Federico II, P.le Tecchio 80, 80125 Naples, Italy.

**Supplementary Movie 1:** Z-stack reconstruction of confocal images of fibroblast cultures after 12 days under different conditions (uninduced, *in vitro* myCAFs, IL1α-only induced fibroblasts and *in vitro* iCAFs). Images were taken with the LSM880 (Zeiis) microscope. Nuclei (dapi) is represented in blue and actin (Phalloidin) in green.

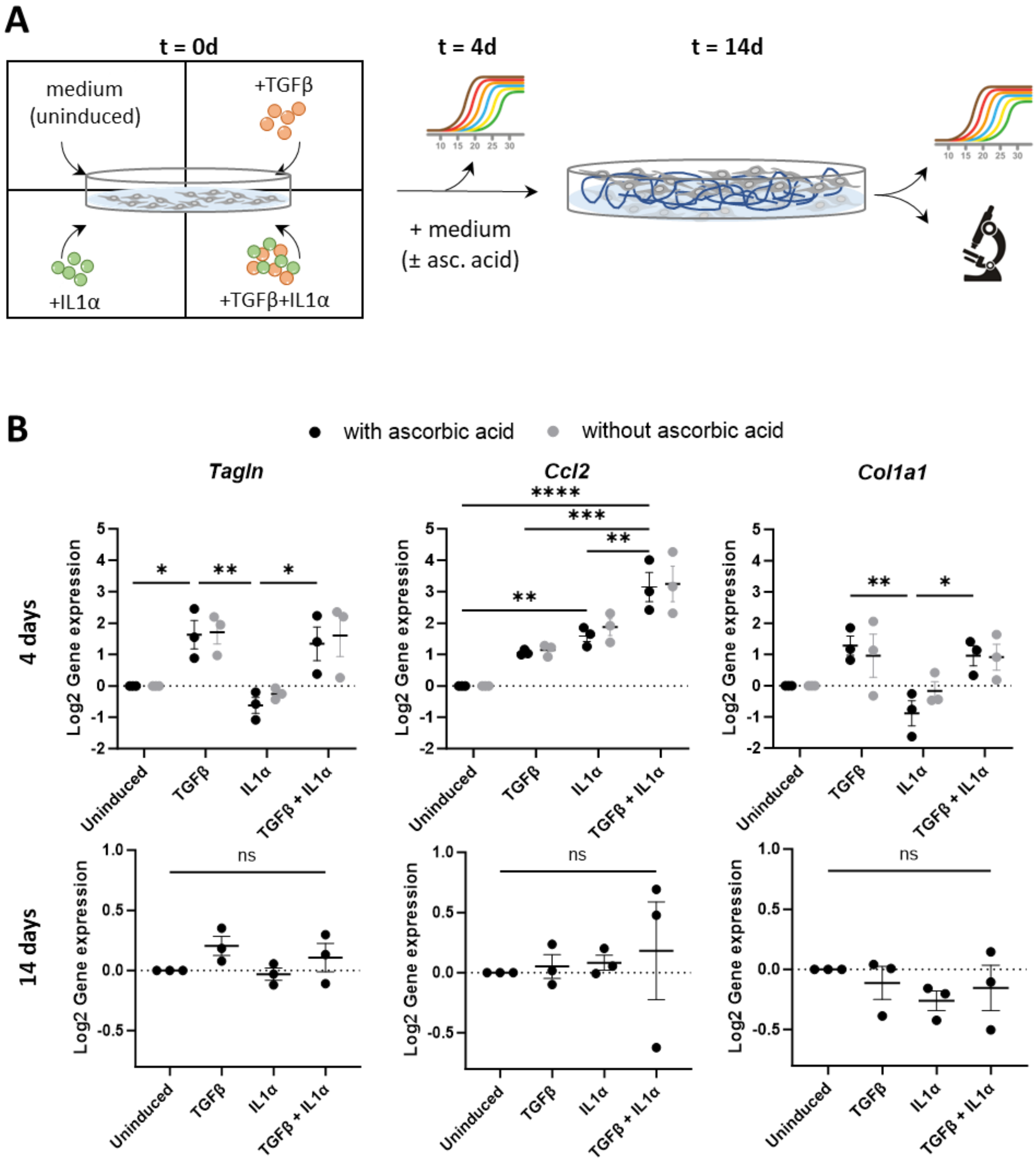


**Supplementary Figure 1: Addition of ascorbic acid does not influence fibroblast phenotype in prolonged cultures to generate cellularized fibroblast subtype-derived matrices. A)** Scheme of the generation of fibroblast subtype specific cellularized matrices. **B)** Log2 gene expression of *Tagln*, *Ccl2* and *Col1a1* after 4 and 14 days normalized to uninduced. Housekeeping genes are *Ppia* and *B2m*. Data were analysed by two-way ANOVA for 4 days gene expression and one-way ANOVA for 14 days gene expression, all followed by Tukey's multiple comparisons test of three independent experiments. *p<0.05; **p<0.01; ***p<0.001; ****p<0.0001.
